# Millisecond cryo-trapping by the *spitrobot* crystal plunger simplifies time-resolved crystallography

**DOI:** 10.1101/2022.09.20.508674

**Authors:** Pedram Mehrabi, Sihyun Sung, David von Stetten, Andreas Prester, Caitlin E. Hatton, Stephan Kleine-Döpke, Alexander Berkes, Gargi Gore, Jan-Philipp Leimkohl, Hendrik Schikora, Martin Kollewe, Holger Rohde, Matthias Wilmanns, Friedjof Tellkamp, Eike C. Schulz

## Abstract

We introduce the *spitrobot*, a protein crystal plunger, enabling reaction quenching via cryo-trapping with millisecond time-resolution. Canonical micromesh loops are mounted on an electropneumatic piston, reactions are initiated via the liquid application method (LAMA), and finally intermediate states are cryo-trapped in liquid nitrogen. We demonstrate binding of several ligands in microcrystals of three enzymes, and trapping of reaction intermediates and conformational changes in macroscopic crystals of tryptophan synthase.

---

Time-resolved X-ray crystallography helps to understand how biomolecular structures change as they carry out their work, offering structural insight into *out-of-equilibrium* conformations and reaction intermediates that cannot be provided by other methods ^1^. Typically, a reaction is initiated in a protein crystal, which is subsequently exposed to an X-ray pulse after a defined delay time. These temporal snapshots can then be assembled into ‘movies’, providing insight even into the fastest processes of life ^2^. However, the majority of enzymes display only moderate turnover kinetics of ∼1 s^-1^, making them accessible by synchrotron radiation experiments^3^. Primarily aiming for these biologically relevant time scales, we have recently developed the *hit-and-return* (HARE) and the liquid application method for time-resolved analyses (LAMA) for *in-situ* mixing ^4,5^. The versatility of these methods encouraged us to further facilitate time-resolved experiments and make them more accessible to the large user base that has already access to standardised tools for high-throughput macromolecular crystallography at synchrotron beamlines.

To this end, we have developed the *spitrobot* crystal plunger, which enables cryo-trapping experiments with versatile time-resolutions down to the millisecond range via the LAMA method (**Figure 1**). Conceptually similar to cryo-EM vitrification devices ^6^, the *spitrobot* relies on crystals mounted onto SPINE-standard micromesh loops (**Supplemental Information**). To trap reaction intermediates the micromeshes are mounted on an electropneumatic piston in an environment with controlled humidity and temperature (**Supplemental Information**). A sequence of electronic signals initiates the *in-situ* mixing reaction by shooting a burst of picoliter-sized droplets onto the mesh-mounted crystals using our established LAMA technology^7^ (**Supplemental Information**). After a defined delay time, the micromeshes are directly plunged into SPINE-standard pucks submerged in liquid nitrogen. As a quality control, sample images are automatically acquired before and after droplet deposition. Adhering to the SPINE standard simplifies the integration into established high-throughput beamline workflows.Characterisation of the processes demonstrate (i) a vitrification time of ∼7.5 ms, for samples on the order of 10 µm, (ii) that physiological conditions can be maintained within the sample area and (iii) that trapping of intermediates can be achieved within 50 ms (**Figure 1, Supplementary information**).

**Figure 1:**
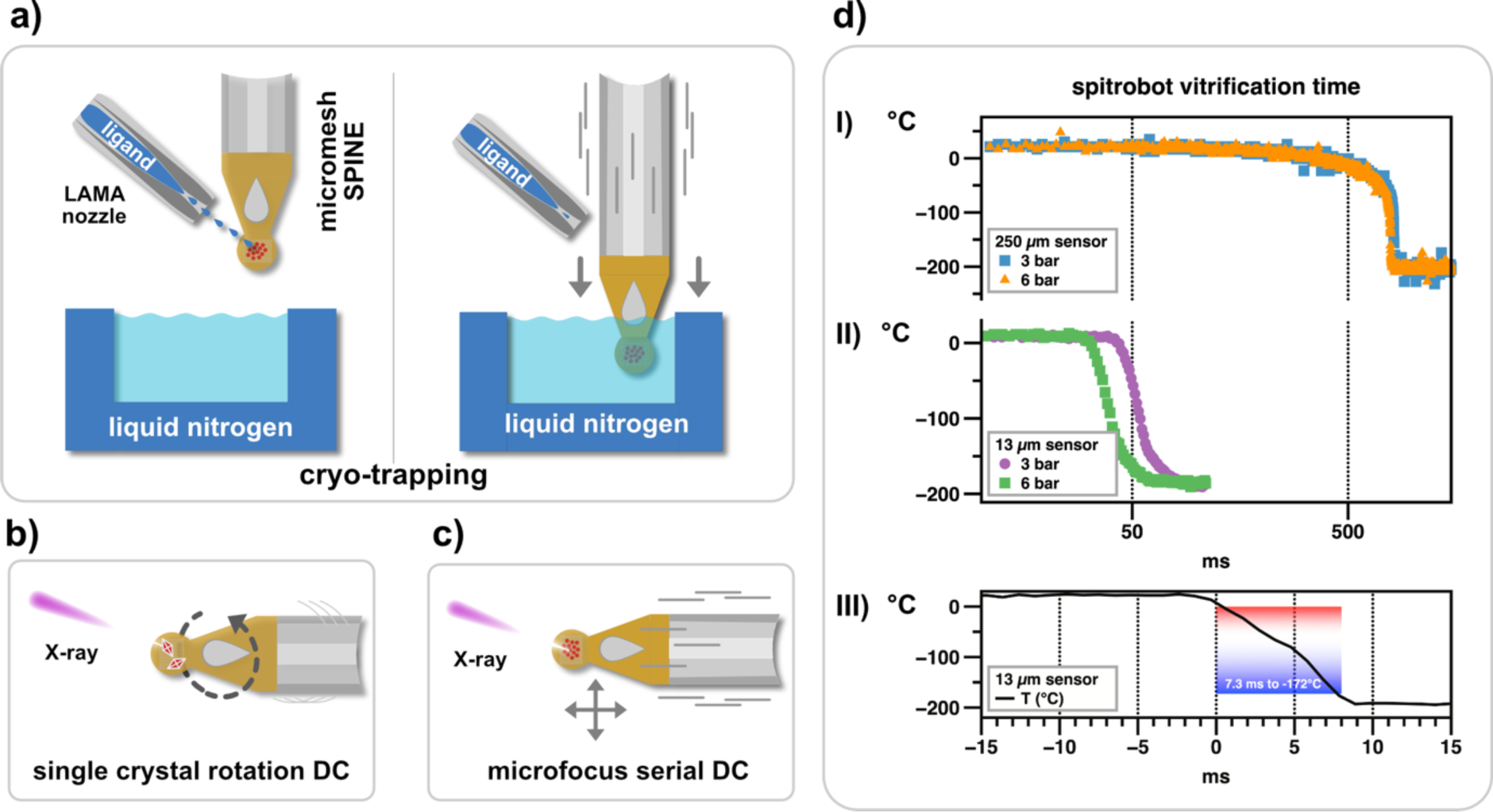
Working principle of the ‘spitrobot’ and characterisation of the vitrification time. (a) crystals are deposited onto micro-meshes; a reaction is initiated via the LAMA method; after a defined delay time reaction intermediates are vitrified in liquid nitrogen. (b,c) the spitrobot integrates with high-throughput workflows and enables using macroscopic crystals and microcrystals for canonical rotation as well as cryo-SSX data collections (d) I,II) vitrification delay determined with 15 µm temperature sensors matching the sizes of typical samples III) an experimental characterisation of the vitrification time demonstrates that the glass transition temperature (160 K) of a typically-sized sample is reached within ∼6.5 ms. The total minimal vitrification delay for microcrystalline samples is approximately 50 ms.

As a first *proof-of-principle* we focused on serial synchrotron data-collection (cryo-SSX). For optimal comparison we used *Streptomyces rubiginosus* xylose isomerase (XI) and the *Klebsiella pneumoniae* extended spectrum β-lactamase CTX-M-14 as model-systems, determining a ligand complex 50 ms and a covalent complex 1 s after reaction initiation, respectively (**Figure 2, Supplementary information**). Next, we explored the potential of canonical rotation data-collection from single crystals prepared with the *spitrobot*. To this end we determined the acyl-enzyme complex of the activity impaired CTX-M-14 mutant E166A in complex with ampicillin, 0.5 s, 1 s, and 5 s after reaction initiation, as well as XI glucose complexes 50 ms, 250 ms, and 1 s after reaction initiation. This emphasizes the high reproducibility of the results, the comparability to TR-SSX data and the suitability for time-resolved experiments in the sub-second time-domain (**Figure 2**).

**Figure 2:**
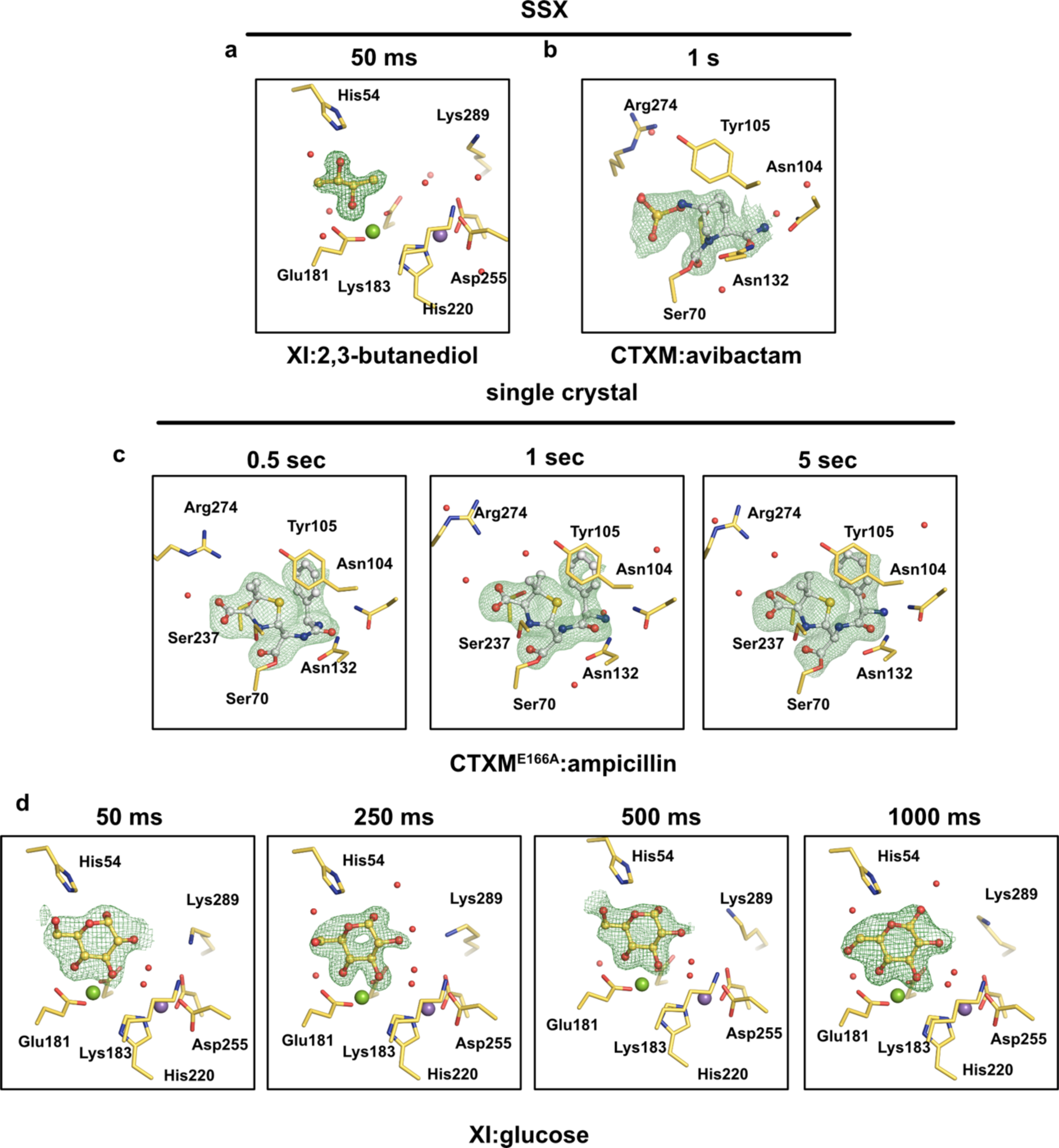
Crystallographic assessment of representative time points. (**a**,**b**) **cryo-SSX**: XI 2,3-butanediol complex formation after 50 ms and CTX-M-14:avibactam complex formation after 1 s. **(c**,**d) single crystals:** CTX-M-14E166A:ampicillin complexes after 0.5, 1 and 5 s, respectively and XI:glucose complexes after 50ms, 250ms, 500 ms and 1000 ms after reaction initiation. POLDER omit maps are shown at 3.0 r.m.s.d.

Finally, we aimed to demonstrate that the *spitrobot* can trap enzymatic reaction intermediates using single, macroscopic crystals. To this end we used the *Salmonella typhimurium* tryptophan synthase (TS), which catalyzes the final steps in tryptophan biosynthesis. We followed the catalytic cycle of its β-subunit, obtaining four structures at 0 s, 20 s, 25 s and 30 s after reaction initiation. The structure at 25 s visualizes the start of the β-subunit reaction. Here serine approaches the internal aldimine, priming the β-subunit for the formation of the external aldimine (Aex-Ser), which can be clearly observed at 20 s and 30 s. Thus, the selected time-points presumably show snapshots of different cycles of the irreversible turnover reaction of the TS β-subunit. Corroborating the turnover, the stabilization of the TrpA-loop6 can be observed in the enzyme 30s after reaction initiation (**Figure 3**). These results demonstrate that the *spitrobot* can conveniently trap reaction intermediates in macroscopic crystals providing insight into enzymatic turnover, at time-scales inaccessible to manual procedures.

**Figure 3:**
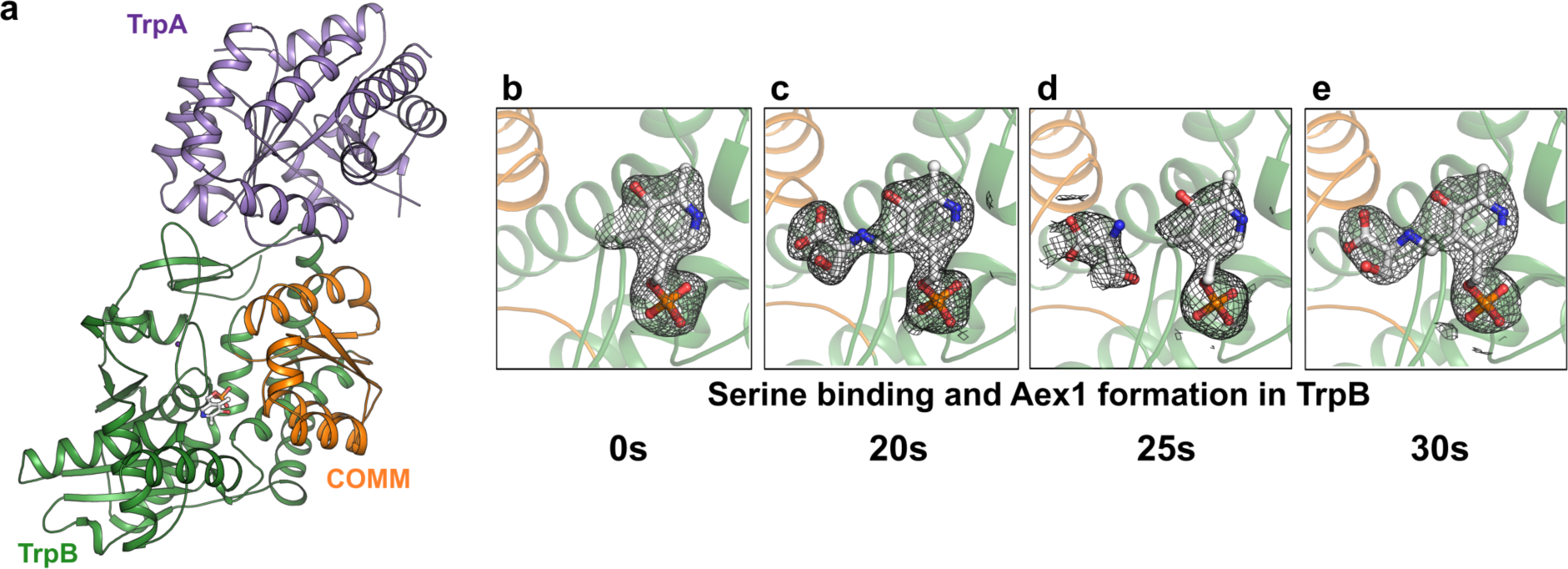
Time-resolved analysis of Tryptophan synthase (TS) turnover. Cartoon representation of Tryptophan synthase AB complex with each subunit represented in a different color. **b-e)** Formation of an external aldimine intermediate (Aex1) at different time points (20 s, 30 s) and serine binding (25 s) after mixing. All electron density maps are represented as composite omit maps contoured at 1.0.

Most available plungers are mainly intended to vitrify cryo-EM samples^8^. By contrast, the commercial plunger ‘*Nanuq’* (Mitegen, USA) is specifically designed to vitrify protein crystals but can -*to the best of our knowledge*-not initiate reactions. More recently a similar crystal cryo-trapping solution was reported ^9^. In contrast to the *spitrobot*, this ‘mix-and-quench’ device relies on non-standard crystal loops without bases and thus requires more complex crystal handling under cryo-conditions, which bears the potential for errors. Moreover, as the crystals are plunged through a substrate-containing film at high velocity it appears difficult to realise the biologically important long-time delays, which greatly limits its applicability to macroscopic crystals. By contrast, the *spitrobot* is built around the SPINE-standard, and versatile time-delays have been demonstrated with both microscopic and macroscopic crystals.While true time-resolved crystallographic experiments at room temperature offer a much more complete access to the dynamic landscape of protein function, cryo-trapping experiments may be sufficient to solve many important biologically relevant questions – e.g. provide insight into thermodynamically trapped, stable reaction intermediates^10,11^. Hence, *spitrobot* experiments provide a number of clear and important advantages: (i) remote time-resolved experiments by uncoupling of sample preparation from data-collection; (ii) makes sub-second cryo-trapping possible, which is manually impossible, and produces longer time delays more consistently than manual harvesting; (iii) preparation for true time-resolved experiments with “a few” crystals to test, e.g., *in-situ* mixing and investigate *in-crystal* kinetics; (iv) enabling systems with an unfavorable crystal-size to diffraction ratio, unsuitable for serial crystallography; (v) physiological conditions during reaction initiation; (vi) enabling low-brightness beamlines or home-sources to carry out time-resolved experiments, as measuring cryo-trapped intermediates is not limited by the photon flux; (vii) most importantly, full compatibility with SPINE standards and thus with the existing high-throughput infrastructure, available at most MX-beamlines. While caution must be applied to mechanistic conclusions that could be biased by the cryo-trapping process (e.g., side-chain conformations, hydration structures, etc.) the practical advantages and its widespread applicability put the *spitrobot* into an ideal position for the transition from static structure determination to real time-resolved crystallography projects^12^. The versatility of the *spitrobot* (crystal size, data-collection routines, time-delays, environmental control), provides ample target opportunities for a large number of labs and beamlines. Importantly the simplicity of the workflow including canonical data-processing, makes it also accessible to the inexperienced users.

## Acknowledgements

Data were collected at beamlines P13 and P14 operated by EMBL Hamburg at the PETRA III storage ring (DESY, Hamburg, Germany). We would like to thank our colleagues A.R. Pearson and G. Bourenkov for helpful discussions and critical reading of the manuscript.

## Author Contributions

E.C.S. designed the experiment.

E.C.S., P. M. and F.T. designed the *spitrobot*.

E.C. S., P. M. and S.S. performed the experiments with support from D.v.S.

P.M., A.P., A.B. and S.S. prepared the protein crystals.

E.C.S., P.M., S.S. and D.v.S. processed and analyzed the diffraction data.

C.E.H., S.K-D., S.S. and G.G. refined crystal structures

JP.L., H.S. M.K. and F.T. built the *spitrobot*, the humidity flow device (HFD) and designed the LabView interfaces.

P.M. and E.C.S. wrote the manuscript.

All authors discussed and corrected the manuscript.

## Online Methods

### Protein crystallization

#### CTX-M-14

CTX-M-14 crystals were generated as described previously^13^. Briefly: Purified CTX-M-14 was concentrated to 26 mg/ml and incubated with CTX-M crystallization buffer (40% PEG8000, 200 mM LiSO_4_, 100 mM NaOAc, pH 4.5) and a highly concentrated seed stock in a 50:45:5 ratio for batch micro-crystallization of the protein. Homogenous micro-crystals with a typical size of 20×20×20 µm were obtained within one day. Crystals of the activity-impaired E166A mutant were generated under the same conditions.

#### Xylose isomerase

Macroscopic crystals of xylose isomerase (XI) were grown via the sitting drop vapor diffusion method. XI was concentrated to 40 mg/ml and equal volumes of protein solution and XI-crystallization buffer (31% (w/v) PEG 3350, 0.2M LiSO_4_, 0.01M Hepes pH 7.5), were incubated for 4 days until first crystals formed, which were harvested after several weeks at a size of approximately 300 µm in diameter. XI microcrystals were generated as described previously ^4^. Briefly: Purified XI was concentrated to 80 mg/ml and incubated with equal amounts of XI crystallization buffer (35% (w/v) PEG3350, 0.2 M LiSO_4_, 0.01M Hepes pH 7.5). The solution was subjected to vacuum evaporation in a ‘speedvac’-micro centrifuge (Eppendorf, Hamburg, Germany) for 15-20 minutes, yielding homogeneous micro-crystals with dimension of 10×15×15 µm.

#### Tryptophan synthase (TS)

TrpA and TrpB were purified as described previously^14^. The TS complex crystallized in 17 % (w/v) PEG 300, 0.1 M Tris-HCl (pH 7.5), and 20 mM Cesium chloride. Crystals grew after mixing 2 µl at 8 – 9 mg/ml protein solution in size-exclusion buffer (50 mM Tris-HCl pH 7.5) with 2 µl of the reservoir solution at 18 °C by the hanging-drop vapor diffusion method. Crystals appeared after 2 – 3 days and reached the final size (0.2 × 0.1 × 0.05 mm) after five to seven additional days.

### Reaction initiation

While microcrystal slurries (∼500 nl) are deposited with a pipette, single-crystals are fished manually and quickly placed in the humidity stream on the *spitrobot*, excess mother liquor is manually blotted via Whatman paper. For reaction initiation the substrate solutions were sterile filtrated and degassed via sonication for 30 minutes. The substrate solutions were loaded into the LAMA nozzles according to the manufacturer’s instructions (Microdrop technologies, Norderstedt, Germany). For complex formation with CTX-M-14 500 150 pl droplets (∼75 nl) of avibactam-buffer: (0.5 M avibactam, 0.14M LiSO_4,_ 0.07 M NaOAc, 0.006 M MES, 15% (v/v) 2,3-butanediol) or 500 droplets of ampicillin-buffer (1 M Na-ampicillin, 0.14M LiSO_4,_ 0.07 M NaOAc, 0.006 M MES, 15% (v/v) 2,3-butanediol), respectively, were applied at a frequency of 2 kHz. For complex formation with XI 200-250 droplets of glucose-buffer (1M Glucose) or 200-250 droplets of butanediol-buffer (1M glucose, 15% (v/v) 2,3-butanediol), respectively, were applied at a frequency of 5 kHz. For reaction initiation with TS, 500 droplets of reaction-buffer (17 % (w/v) PEG 300, 0.1 M Tris-HCl (pH 7.5), 20 mM Cesium chloride, 10 mM G3P, 110 mM indole, 100 mM serine and 30 % ethanol) were applied at a frequency of 6 kHz via the LAMA nozzle.

### Data-collection and processing

#### Cryo SSX

Serial synchrotron crystallography was originally established under cryo-conditions using a limited rotation workflow^15^. However, unlike in the original workflow we collected still diffraction images using a mesh scan workflow available in MXCuBE. A double collimated beam with a FWHM size of 7×3 µm, at an energy of 12.7 keV (0.9763Å) at a flux of ∼2 × 10^13^ ph/s and an exposure time of 7.5 ms per image was used during data-collection with an Eiger2 CdTe 16M detector (Dectris, Switzerland). For the mesh collections, a mesh with a grid spacing matching the dimensions of the beam was drawn over the whole micro-mesh sample, giving rise to several thousand still diffraction images, which were processed using CrystFEL with the XGANDALF indexing routine ^16,17^. Structures were solved by molecular replacement in PHASER using 6GTH as a search model for CTX-M-14 and 6RNF as a search model for XI ^18^.

#### Single crystal data

Cryo-trapping data from single crystals were solved by making use of canonical, single crystal data-collection workflows. A double collimated beam with a FWHM size of 7×3 µm, at an energy of 12.7 keV (0.9763Å) at a flux of ∼4 × 10^11^ ph/s and an exposure time of 7.5 ms per image was used during data-collection on an Eiger2 CdTe 16M detector (Dectris, Switzerland). Diffraction data were processed using XDS^19–21^ and AutoPROC using StarAniso^22,23^. For processing the TS datasets, the collected datasets were initially integrated using XDS and merged and scaled using the CCP4 suite program AIMLESS^24,25^. Structures were solved by molecular replacement in PHASER using 2WSY as a search model for TS^26^, 6RNF as a search model for XI, and using 6GTH as a search model for CTX-M-14.

### Refinement & Data analysis

Refinement was carried out in the phenix suite using *phenix*.*refine* ^27^and coot 0.8 for manual corrections to the model ^28^. POLDER maps were generated using *phenix*.*polder*^29^. Composite omit maps for TS were generated using *phenix*.*composite_omit_map*. Molecular images were generated in PyMol ^30^.

## Supplementary Information

### The *spitrobot*

The *spitrobot* comprises several different, main hardware parts: a) the plunger, b) the humidity flow device (HFD), c) the LAMA droplet injector, d) the vitrification chamber, f) the camera system, and e) the control unit. All parameters are set via a control software.

### The plunger

The main component of the plunger is an electropneumatic piston that drives the sample into the liquid nitrogen. It is mounted on a sturdy steel post on top of the vitrification chamber. The plunging velocity is regulated via the applied gas pressure. For typical use we relied on pressure levels from 3-6 bar, which enable piston motions on the order of 1.6 ms^-1,^ comparable to previously published solutions ^9,31^. The piston is equipped with an electromagnetic SPINE-style sample holder, onto which the micro-meshes are manually mounted. After being submerged in liquid nitrogen the micro-meshes are automatically released into the SPINE-pucks, minimizing manual interaction after sample preparation. For reaction initiation the LAMA nozzle needs to be positioned within ca. 1 mm of the micro-meshes. To avoid accidental collisions and simplify sample mounting, the LAMA nozzle is retracted via rail-mounted translation stages. Once the sample is mounted, the LAMA nozzle is pushed back into place, and subsequently fine-aligned to the micro-mesh. Using the SPINE standard provides a number of advantages regarding the compatibility with established high-throughput beamline workflows. Adhering to this established standard will streamline crystal storage and shipment from (remote) MX-labs to synchrotron facilities, where sample exchange and automated data-collection procedures also rely on standardized samples, greatly increasing the turnover of samples and even enabling fully automated data-collection^32^.

**Supplementary Figure 1:**
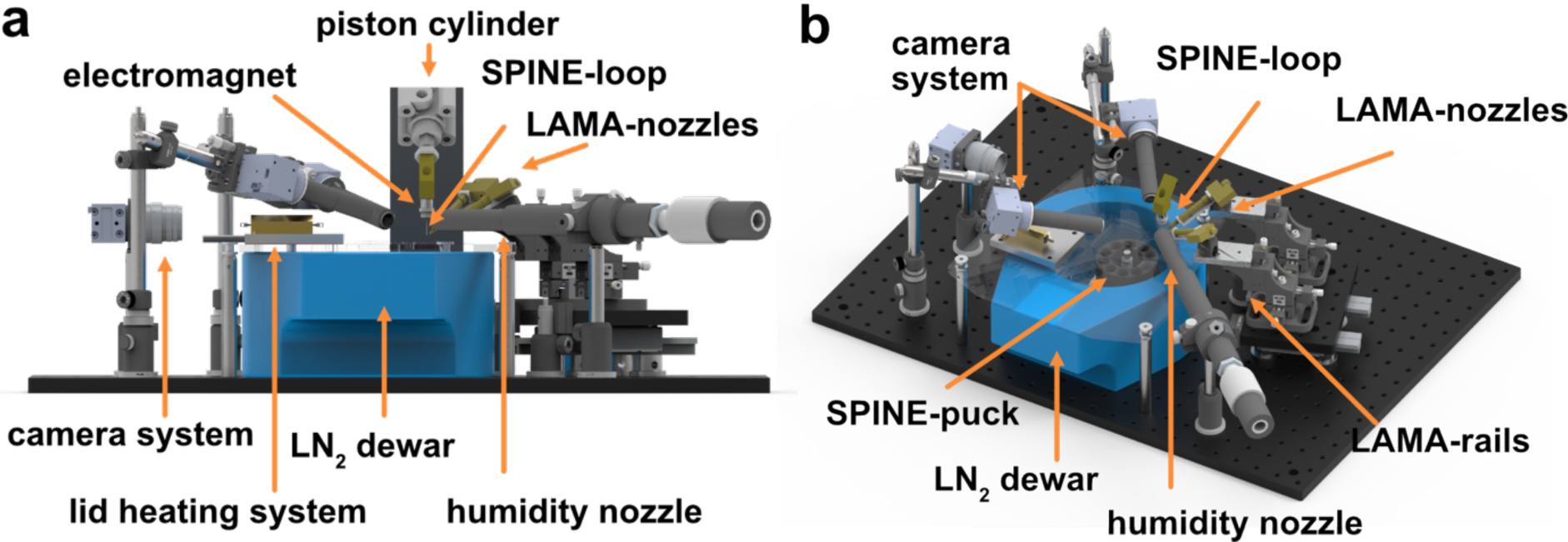
The experimental setup of the spitrobot plunger. a) front view b) isometric view, the piston cylinder has been omitted for clarity. Relevant parts are labelled, please refer to the text for further details.

### Environmental control with the HFD

To maintain the crystals in an environment close to physiological conditions with controlled humidity and temperature we have developed a Humidity Flow Device (HFD), which provides a humid airflow at a defined temperature between 4 °C and 40 °C (**Supplementary Figure 2**). Both the relative humidity and the temperature can be adjusted independently. The nozzle of the HFD has an inner diameter of about 13 mm and is placed about 1 cm from the SPINE sample. On the inside of the nozzle, a few cm before its end, is a combined sensor for humidity and temperature whose signals are transmitted to a microcontroller (Arduino nano), which calculates the control variables. The temperature is controlled by heating resistors; the humidity is controlled by a warm water bath, which is equipped with ultrasonic nebulizers. An external cooler can also be connected via a heat exchanger. The HFD provides temperatures between 4 °C and 40 °C at a humidity of up to 99%, with typical flow rates between 20 and 35 lpm. Humidity control also permits controlled crystal dehydration if that is required (see separate section below). To characterise the stability of the HFD we recorded step-functions of the relative humidity and the temperature, respectively, as a function of time. Humidity was increased at 5% increments and maintained at stable flow for several minutes. After an equilibration period, the relative humidity can be maintained within less than a percent (**Supplementary Figure 3**), demonstrating its suitability to maintain a stable humidity environment for crystals and micro-crystals during sample preparation (**Supplementary Figure 4a**). For the temperature step function, the temperature was recorded at 4°C, 10°C, 20°C, 30°C and 40°C for several minutes, respectively, keeping the relative humidity constantly at or above 95%.

**Supplementary Figure 2:**
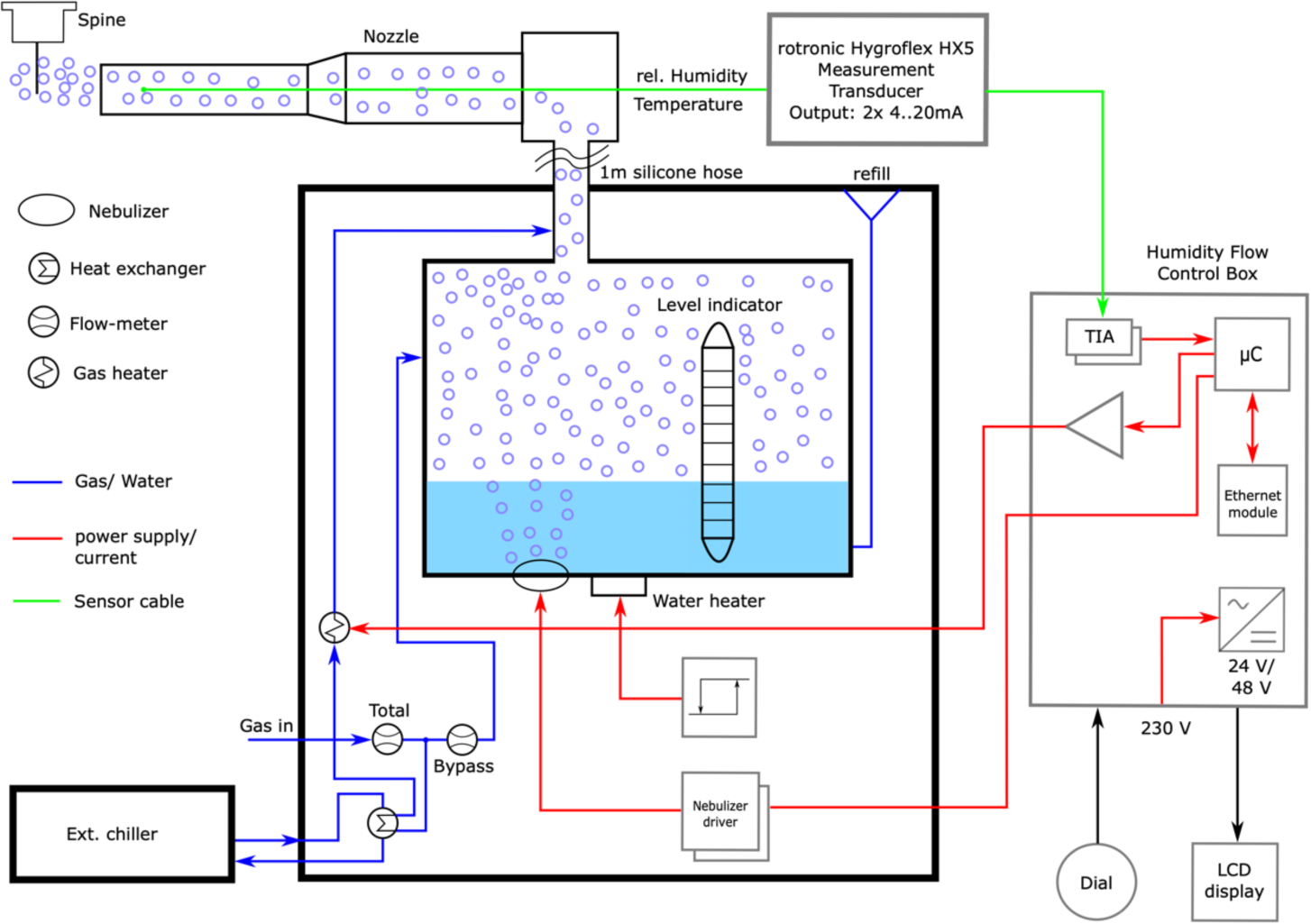
Schematic overview of the humidity flow device (HFD). The scheme shows the interplay of the various hardware parts of the HFD.

**Supplementary Figure 3.**
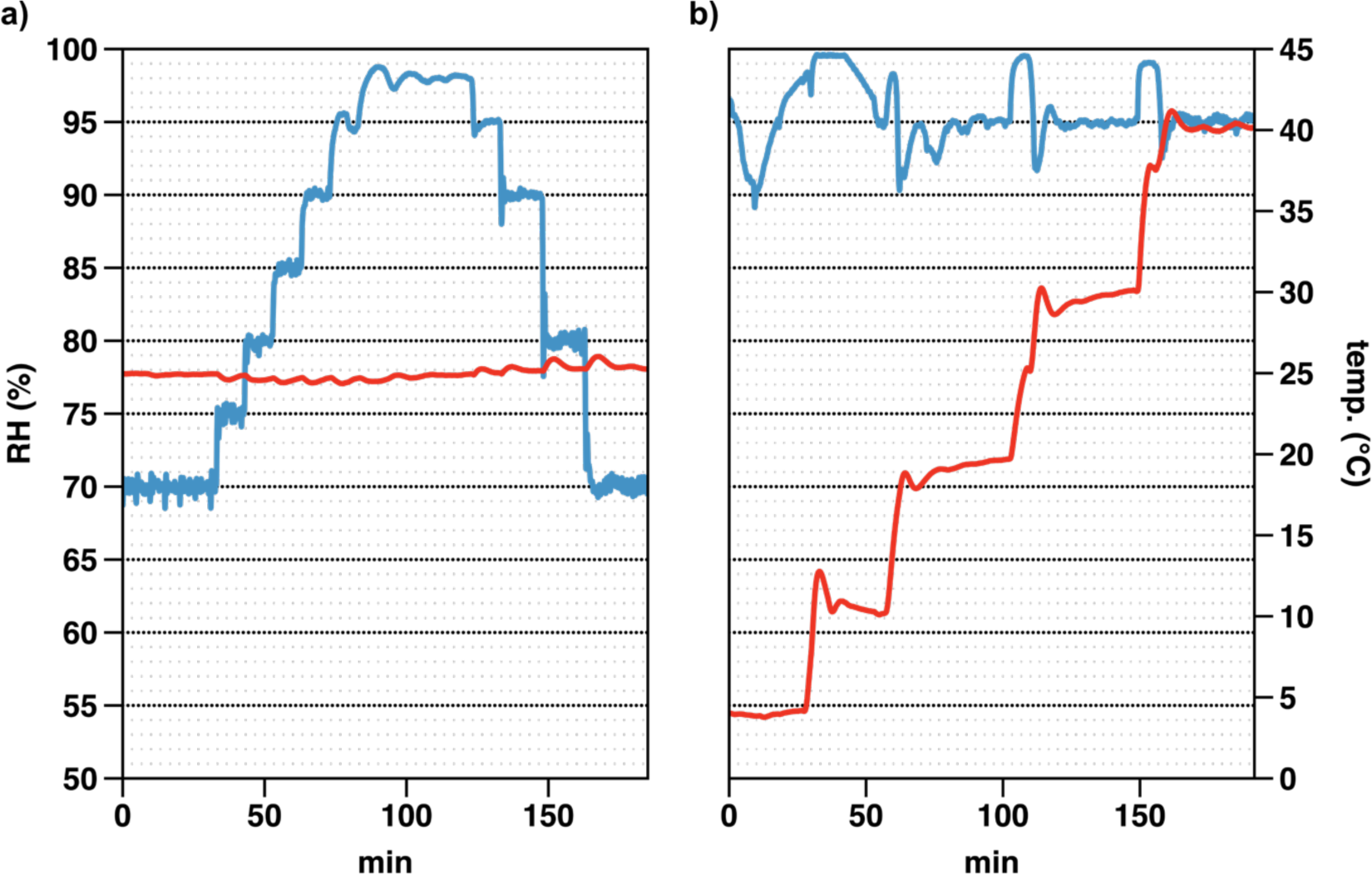
Characterisation of the HFD. Environmental parameters are displayed as a function of time. **a**) a step-function of the relative humidity at constant temperature (25°C). **b**) a step-function of the temperature at constant relative humidity (95%).

### Reaction initiation

The LAMA droplet injector has been previously described in detail ^4^. Briefly: via a piezo-actuator, 75 or 150 picoliter sized droplets are shot from a 50 or 70 µm glass capillary with a velocity of 2 m/s onto the target mesh. The nozzle is brought close (1-2 mm) to the target via manual, rail-mounted translation stages that enable precise lateral and vertical alignment of the nozzles to adjust for differences in SPINE loop length and also correct for, e.g., bent loops. Nozzle alignment is aided by two perpendicularly aligned cameras, which focus on the target mesh (see below). This way the nozzle distance, as well as its lateral and vertical alignment can be precisely controlled and adjusted to individual loops. Since the micromeshes involve a large sample area, a high-frequency (5 kHz) burst of picolitre droplets are added to the samples. The total volume of required liquid depends on the sample area that has to be covered, the protein and ligand concentrations, and the viscosity of the solutions. Between 100 and 500 droplets were applied for each sample used in this study.

### The vitrification chamber

The vitrification chamber is comprised of a regular foam dewar into which an aluminum mount for the SPINE standard puck is fixed. The mount permits a step-wise rotation of the puck between its 10 positions for sample vials for aligning each position to the vitrification point of the *spitrobot*. Rotation of the puck to the next sample position is done manually via a hexagonal bolt screw driver. This simplifies and accelerates sample handling and transfer as the process of vitrification deposits the sample directly into the puck, enabling further usage of high-throughput infrastructure. The dewar is closed via a transparent poly(methyl methacrylate) (PMMA) lid, with an opening for the piston. To reduce icing in the liquid nitrogen phase and improve vitrification rates the gaseous nitrogen layer that forms between the lid and the liquid phase is displaced by a stream of dry nitrogen gas as demonstrated previously^1^. To this end the dry nitrogen is actively siphoned away via a connected pump. To further reduce ice formation on the lid, it is heated via a resistor array. Liquid nitrogen is replaced manually at regular intervals via the refill hole or the piston opening.

### Experimental characterisation of the vitrification process

The vitrification time was characterised by two independent approaches, optically and electronically. For an optical characterisation an LED flashing every 2.5 ms (400 Hz) was mounted to the tip of the piston. A long time-exposure synchronized to the plunging process captured the number of flashes during the piston motion. Since 9 flashes were recorded, this is equivalent to a piston motion time of 22.5 ms. However, this procedure only characterises the piston motion, thus the switching delay of the air valve and the actual vitrification time were addressed in a separate experiment.

To obtain accurate vitrification times we used a thermocouple within similar dimensions to the crystal size. For comparison to previous studies, which mainly used larger thermocouples we thus recorded vitrification times using two temperature sensors of different size. The larger RTD (3.0 × 0.8 × 0.25 mm (IST, P1K0.308.7W.B.007 Farnell, Germany, -200°C – 600°C) displays a total time to quench of 800 ms, which is presumably offset due to the Leidenfrost effect. By contrast, the smaller thermocouple, which has approximately the same dimensions as the samples of interest (K-type thermocouple [KFT-13-200-200(Y)], ∼13 µm diameter, ANBE SMT Co., Osaka, Japan) minimizes this offset and displays a total time to quench of ∼50 ms and is thus almost negligible for the relevant time scales (**Figure 1d**). To determine the actual vitrification time, we also recorded the temperature decrease independent of the *spitrobot*. Here the glass-transition temperature (< -140 °C) is reached within 6.5 ms and a 90-10% analysis of the temperature drop resulted in a fall time of 7.5 ms. Thus, the total delay time of the *spitrobot* consists of the intrinsic delay of the device (air valve delay ∼20 ms, piston motion ∼25 ms), and the actual vitrification process ∼7.5 ms corresponding to a cooling rate of 2.3×10^4^ Ks^-1^. These vitrification times are comparable to those reported previously for flash cooling devices operating with liquid nitrogen^31^. Based on these observations the dead-time of the *spitrobot* is on the order of ∼45 ms and time-points with a minimal delay time of approximately 50 ms can be obtained.

### Camera system

For automatic reference image acquisition and convenient sample alignment, the *spitrobot* is equipped with two cameras from different viewing directions, 90° apart. The primary purpose of the camera system is sample alignment and LAMA-nozzle positioning. The axis of the LAMA nozzle is at an angle of 60° relative to the surface of the mesh. The LAMA nozzle can be positioned using the aforementioned 3-axis stage to optimally deposit the substrate on the micromesh and ensure reproducible results between different samples. By using the high-resolution camera system during sample preparation, the consistency between different samples can be maintained. In addition, this setup provides automatic image acquisition immediately before and after droplet depositions. This serves as a reference to a) confirm the droplet depositions and b) droplet dissipation on the micromesh. The latter can be used during data collection at the beamline to narrow down the area for data collection (**Supplementary Figure 4**).

**Supplementary Figure 4:**
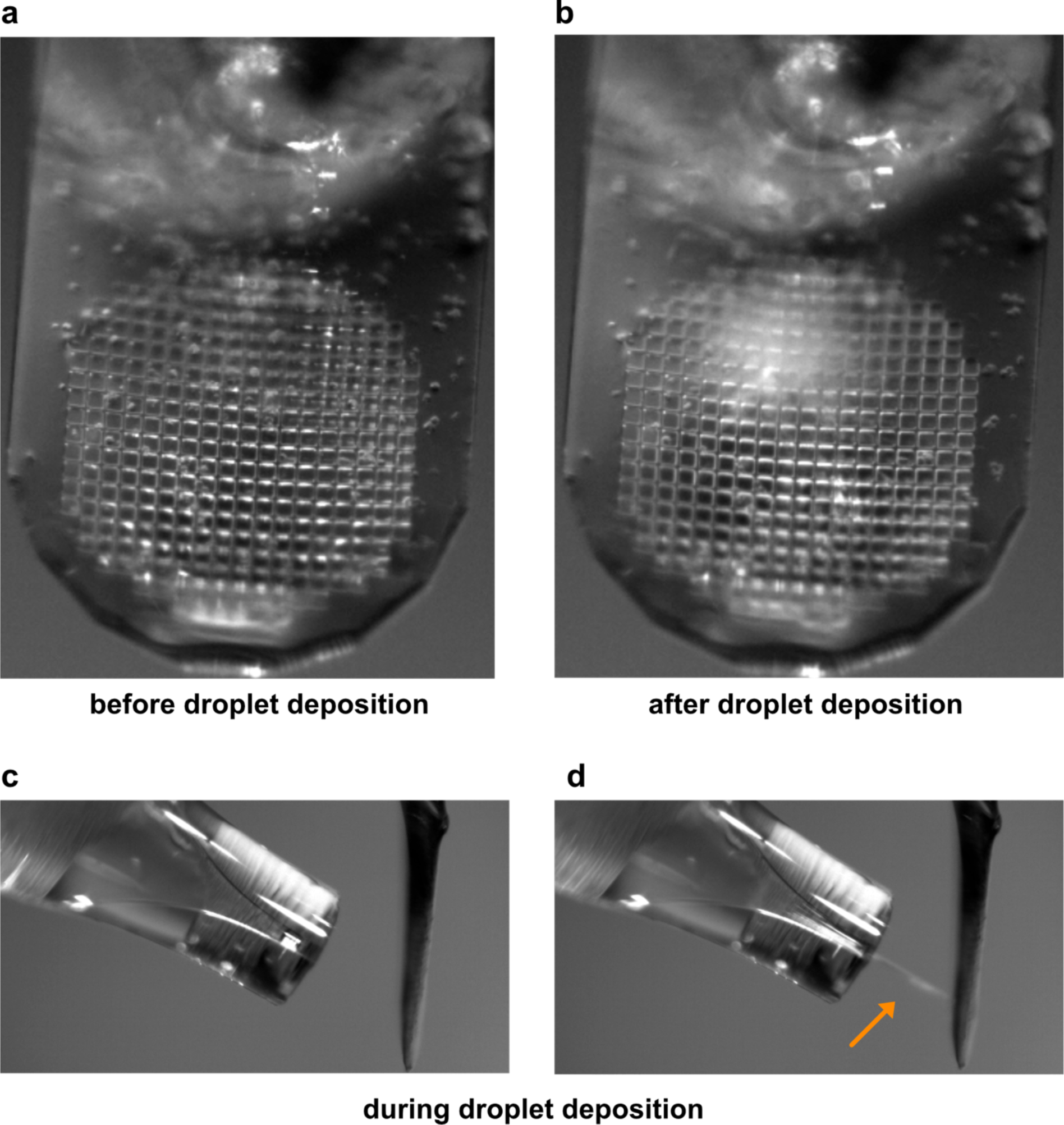
Automatic assessment of droplet deposition. **a-b)** The figures show a comparison of a typical micro-mesh loaded with crystals a) before droplet deposition, and b) after droplet deposition. The nozzle outlet is visible in the top part of the figure, behind the semi-transparent micro-mesh material. The side view (c,d) shows the nozzle to micro-mesh distance and the droplets in flight.

### Control unit and control software

The individual steps of this protocol (reference image acquisition, droplet deposition (reaction initiation), control image acquisition, and piston motion after the pre-defined delay time) are triggered by a two-hand control device (THCD) integrated into the control unit. The two-button operation is a mandatory safety requirement to prevent inadvertent interaction with the piston during its motion, which due to its high velocity has the potential to cause serious injuries.

The delay-time settings, as well as automatic image acquisition is realized via a LabVIEW control software. Users can conveniently define file names and storage directories to obtain sample images before and after droplet deposition for quality control and referencing. In addition, users can define the delay time between droplet deposition and vitrification, which is transferred to the safety switch device. The delay time defined by the users is the total delay time for the whole period until the reaction is quenched, i.e., the time for piston motion is taken into consideration.

**Supplementary Figure 5:**
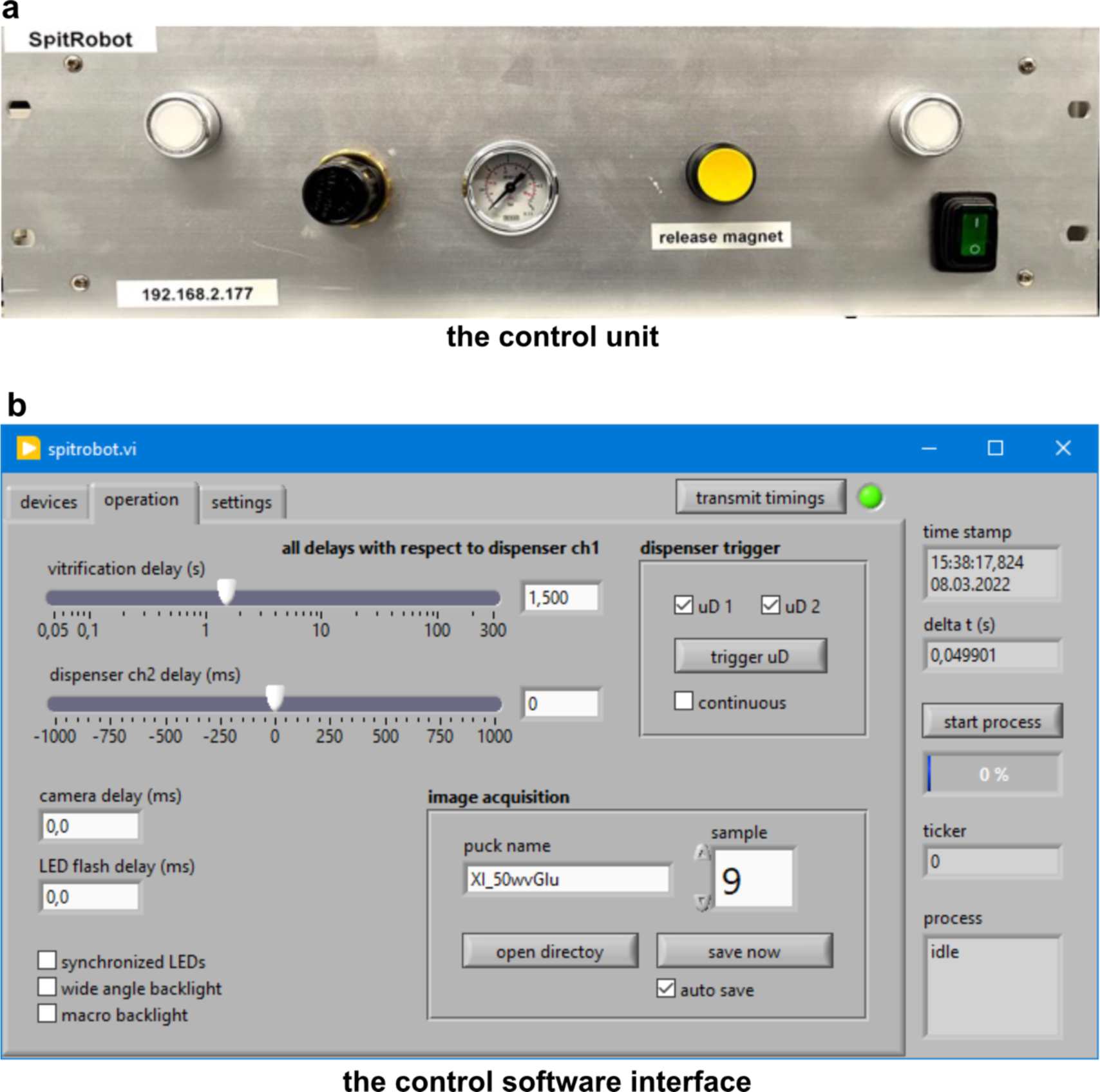
Screenshot of the control software written in LabView. A) The control unit is powered on via the main switch, enables to set the plunging pressure, to release the SPINE sample via an electromagnet and to start the operation cycle by simultaneously pressing the two white trigger buttons. b) via the control software all relevant parameters such as vitrification and delay times, file name and sample numbering can be set and transferred to the control unit.

## Workflow

### *Spitrobot* preparation

The HFD should be switched on approximately 1 h before starting to load samples to allow for equilibration of temperature and humidity. Immediately prior to mounting in the *spitrobot*, the micromeshes are glow-discharged for 60 seconds in a glow-discharge machine, typically used for EM-grid preparation (CTA 010, Balzers Union, Switzerland). A SPINE crystal storage puck and the vitrification dewar are pre-cooled in liquid nitrogen and then transferred to the *spitrobot*. The LAMA nozzle is loaded with the ligand solution of interest (pre-mixed with an appropriate cryo-protectant if required). Droplet formation conditions must be established by adjusting voltage and pulse width in the Mircodrop controller (Microdrop technologies, Norderstedt, Germany). Due to the individual ligand-solution parameters, droplet distribution on a micromesh and the number of droplets must be established on a dummy loop using the camera video for visual verification. The number of droplets is set in the Microdrop controller.

### Operation

Initially project directory and sample names are defined in the control software, and the desired time delay is set and transferred to the control unit. For crystal loading the LAMA nozzle is retracted to allow better loop access. A micromesh loop is placed on the electromagnet to adjust camera height, position, and focus. Crystals can be loaded either by manual fishing, which works for both macroscopic crystals as well as micro-crystal slurries. Alternatively, micro-crystal slurries can be loaded via droplet deposition from a micro-pipette directly on the micromesh placed in the humidity stream. Next, an approximate nozzle position is adjusted using the manual translation stages. Using the THCD, user safety is maintained and the automated part of the *spitrobot* operation is started: (1) the predefined number of droplets is shot onto the micromesh, (2) after the pre-defined delay time the piston drives the loop into the liquid nitrogen where the crystal is vitrified directly in a precooled vial in the storage puck, (3) after 2 s the electromagnet releases electromagnet releases the loop, and (4) the piston moves back into its starting position. Then the user manually rotates the storage puck into the next position to prepare the system for the next round of operation. Due this partially automated process, the complete loading of a puck with 10 samples takes only about 30 - 60 min.

## Controlled crystal dehydration

To demonstrate the precise control of the humidity at the sample position and highlight the potential for controlled crystal dehydration we directly attached the humidity nozzle to the diffractometer at beamline P14 at EMBL Hamburg. We chose xylose isomerase as a model system. A macroscopic crystal of xylose isomerase was mounted in a canonical nylon loop and placed in the humidity stream at 99% relative humidity. Canonical rotation datasets were collected at consecutively lower levels of relative humidity. To mitigate potential radiation damage, the dose was kept as low as reasonably possible at an average diffraction weighted dose of 3.5 kGy per dataset^33,34^. The consecutive datasets show an overall ∼20% change in unit-cell volume accompanying the reduction in relative humidity, eventually resulting in an alternate spacegroup (**Supplementary Figure 6, Supplemenaty Table 1**). This also reflects in the crystal quality parameters, which surprisingly improve after the change in spacegroup, re-emphasizing the potential of controlled crystal dehydration for data-collection. The details of this observation will be discussed elsewhere. Briefly, unit-cell compactions, including xylose isomerase due to controlled crystal dehydration have been reported previously with devices operating within similar relative humidity levels (∼70-99%) ^35–38^.

**Supplementary Figure 6:**
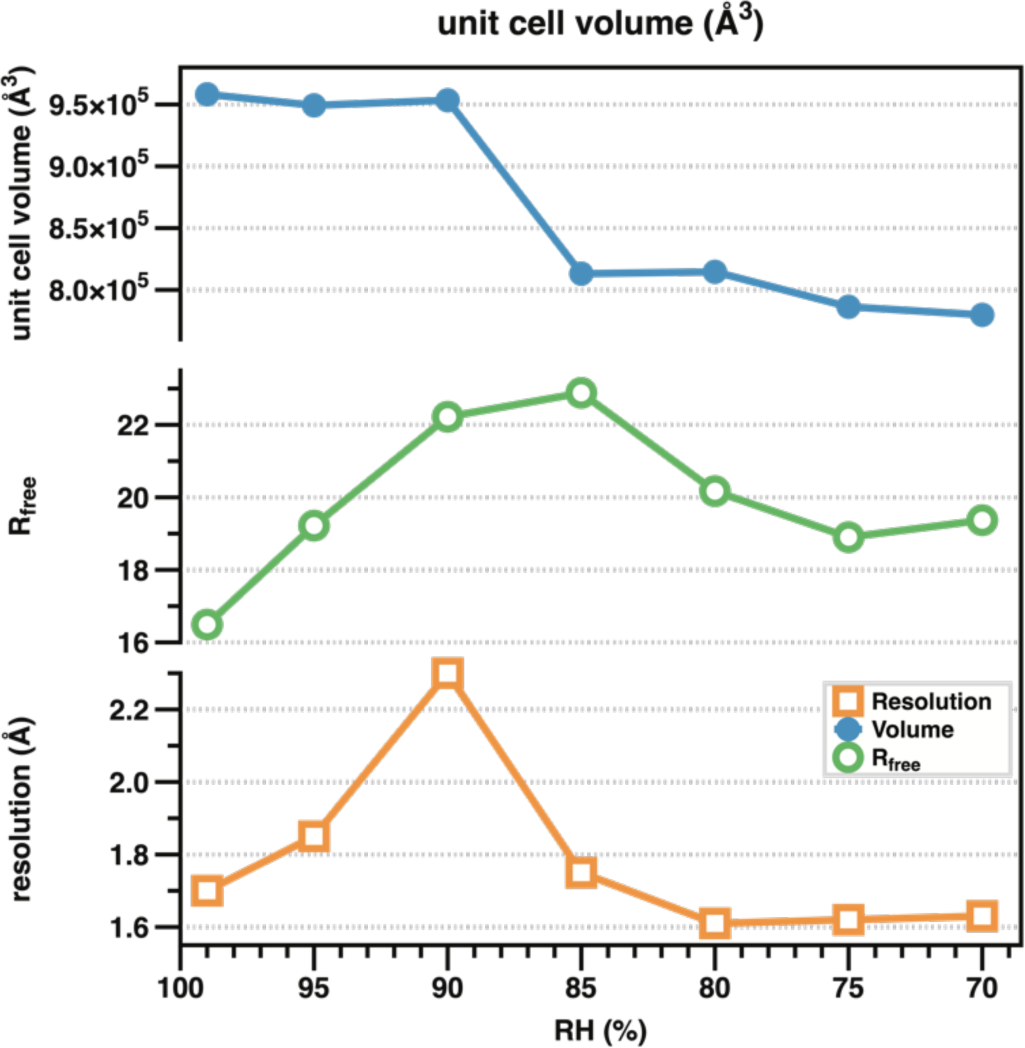
Xylose isomerase unit cell volume and selected data-quality indicators as a function of relative environmental humidity.

**Supplementary Table 1:**

| XI |  |  |  |  |  |  |  |
| --- | --- | --- | --- | --- | --- | --- | --- |
| PDB-ID | 8AWE | 8AWD | 8AWB | 8AWC | 8AWF | 8AW9 | 8AW8 |
| Relative humidity | 99% | 95% | 90% | 85% | 80% | 75% | 70% |
| Resolution | 46.88-1.70 | 49.55-1.85 | 69.14-2.30 | 68.29-1.75 | 43.78-1.61 | 52.99-1.62 | 52.83-1.63 |
| range (Å) | (1.74 - 1.70) | (1.89 - 1.85) | (2.44 - 2.30) | (1.81 - 1.75) | (1.63 - 1.61) | (1.64 - 1.62) | (1.66 - 1.63) |
| R<sub>Free</sub> (%) | 16.49 | 19.22 | 22.22 | 22.88 | 20.17 | 18.90 | 19.37 |
|  | (25.08) | (31.37) | (22.78) | (34.03) | (39.68) | (23.73) | (31.59) |
| R<sub>Work</sub> (%) | 13.58 | 16.00 | 17.38 | 18.98 | 17.87 | 15.47 | 15.98 |
|  | (25.48) | (28.23) | (18.52) | (28.32) | (40.04) | (18.67) | (20.74) |
| Cell | 93.767, | 93.480, | 93.839, | 87.165, | 87.100, | 83.144, | 82.826, |
| Dimensions | 99.397, | 99.110, | 99.357 , | 94.506, | 98.750, | 96.591, | 96.307, |
|  | 102.807 | 102.460 | 102.256 | 98.794 | 94.710 | 97.913 | 97.757 |
| a, b, c (Å); | 90.00, 90.00, | 90.00, 90.00, | 90.00, 90.00, | 90.00, 90.00, | 90.00, | 90.00, | 90.00, 90.00, |
|  | 90.00 | 90.00 | 90.00 | 90.00 | 90.00, | 90.00, | 90.00 |
| α,β,γ (°) |  |  |  |  | 90.00 | 90.00 |  |
| Space Group | I222 | I222 | I222 | I222 | P 21 21 2 | P 21 21 2 | P 21 21 2 |

## Thermodynamic reaction quenching via cryo-trapping

Microcrystals permitting for serial data collection should have minimal dimensions, suitably sized to match the desired microfocus beamline. The dimensions of the crystals also define the minimal ligand diffusion time, which should be faster than turnover time to reduce heterogeneity in the cryogenically trapped states.

### Serial crystallography

To demonstrate that *spitrobotspitrobot*-prepared micro-crystals are suitable for serial cryo data collection, 0.5 µl of the crystal slurry was directly loaded by pipette on the SPINE standard, 700/25 µm micromesh (MiTeGen, USA) and quickly transferred to the humidity stream. Excess mother liquor was manually blotted away until the sample meniscus disappeared, by quickly (< 1 sec) applying Whatman paper to the back of the mesh. For reaction initiation the ligand solution was supplied in the LAMA nozzle, 250-500 droplets were deposited in a burst mode (2-5kHz repetition rate).

For comparison we utilized *Streptomyces rubiginosus* xylose isomerase and the *Klebsiella pneumoniae* extended spectrum serine beta-lactamase CTX-M-14 as model-systems. After a set delay time of 50 ms for xylose isomerase (XI) and 1 s for CTX-M-14 the crystals were vitrified in liquid nitrogen by directly plunging them into a puck. After structure determination clear difference electron density was visible in the active site, which could be interpreted by modelling the ligand molecules (**Figure 2**). Comparison to our previously determined CTX-M-14 structure in complex with avibactam (PDB-ID: 6GTH) reveals only minor differences between the cryogenically cooled crystals and the room-temperature complex determined by SFX^13^. This confirms efficient mixing and diffusion of the ligand into the active site as we have demonstrated previously ^4,13^ (**Figure 2, Supplementary Table 1)**.

Using 2,3-butanediol as a cryo-protectant in a buffer containing its natural ligand glucose, we found that 2,3-butanediol can also occupy the XI active site within 50 ms, and is not replaced by glucose within 500 ms after reaction initiation (**Figure 2, Supplementary Table 1**). The 2,3-butanediol molecule soaked into the XI crystals adopts a conformation similar to our previously determined glucose bound complex structure 15 ms after reaction initiation^4^.

**Supplementary Table 2.** Data collection and refinement statistics for cryo SSX crystal datasets.

|  | **XI** | **CTX-M-14** |
| --- | --- | --- |
| PDB-ID: | 8AWY | 8B3M |
| Delay time | 50 ms | 1000 ms |
| Ligand | 2,3-butanediol | avibactam |
| **Data collection** |  |  |
| Space group | I 2 2 2 | P 3 1 2 1 |
| Cell dimensions (a, b, c (Å)) | 92.7 102.35 99.3 | 41.85 41.85 232.85 |
| $\alpha, \beta, \gamma$ (°) | 90.00 90.00 90.00 | 90.00 90.00 120.00 |
| Resolution (Å) | 71.27 - 1.6 (1.657 - 1.6) | 77.62 - 1.6 (1.66 - 1.60) |
| Number of images | 30137 | 34453 |
| I/$\sigma$(I) | 9.70(5.96) | 6.37 (1.83) |
| CC 1/2 | 97.92(95.19) | 96.37 (73.00) |
| Completeness (%) | 99.97 (99.98) | 100 (100) |
| Redundancy | 662.0(454.3) | 532.7 (369.1) |
| **Refinement** |  |  |
| Resolution (Å) | 1.6 | 28.60-1.95 |
| Total reflections | 41335785 | 32733 |
| Unique reflections | 62424 | 18275 |
| R work / R free | 0.1743/0.1940 | 24.99 / 29.68 |
| **No. atoms** |  |  |
| Protein | 3111 | 1968 |
| Ligand | 9 | 17 |
| Water | 485 | 265 |
| **B factors** |  |  |
| Protein | 13.2 | 24.65 |
| Ligand | 24.16 | 31.44 |
| Water | 27.89 | 30.86 |
| **R.m.s. deviations** |  |  |
| Bond lengths (Å) | 0.005 | 0.006 |
| Bond angles (°) | 0.85 | 0.921 |
| **Ramachandran statistics** |  |  |
| Favored regions (%) | 96.88 | 96.12 |
| Allowed regions (%) | 2.86 | 2.33 |
| Outliers (%) | 0.26 | 1.55 |

## Canonical rotation crystallography

As an alternative to serial crystallography, we aimed to demonstrate that the *spitrobotspitrobot* is also suitable for standard rotation data collection. We used the microfocus beam (3×7 um) of EMBL beamline P14 (Hamburg), where individual micro-crystals were centered in the X-ray beam. A convenient approach to identify well-diffracting crystals is generating a diffractive-power heat map via the mesh collection option in MXCuBE (**Supplementary Figure** 7)^39^. After selection of a suitable crystal a standard rotation dataset was collected, amenable to automatic data-processing routines available at most macro-molecular crystallography beamlines.

### Extended spectrum beta-lactamase CTX-M-14

To demonstrate that such data collections work routinely, we prepared complexes of the activity impaired CTX-M-14 E166A mutant with ampicillin, at time-delays of 0.5 s, 1 s and 5 s after reaction initiation. At all time points the electron density confirms that a covalent acyl-enzyme intermediate has formed, which compares well to previously published data and thus confirms the consistency of the crystallographic data across broad time-scales (7K2Y) (**Figure 2**)^40^. In addition, this experiment demonstrates that cryo-trapping data can successfully be obtained via canonical rotation data collection and automatic data processing routines, which greatly accelerates the structure determination process.

### Xylose isomerase

Next, we aimed to explore the minimal permissible *spitrobot* delay time in comparison to previously established data^4^. For consistency between the results obtained via the LAMA method at room-temperature SSX and the cryo-trapping results from the *spitrobot* we made use of our previously established model system xylose isomerase (XI). XI microcrystals were loaded onto SPINE standard, 700/25 µm micromeshes (MiTeGen, USA), directly inside the humidity stream using a standard micro-pipette. For reaction initiation the substrate solution (1M D-glucose (aq)) was supplied in the LAMA nozzle, 250 droplets were deposited using the burst mode (5 kHz repetition rate).

Previously we had determined by RT-SSX that near full ligand occupancy can be obtained in XI within 15 ms, exceeding what is mechanically feasible with the *spitrobot*^4^. Thus, crystals were vitrified after 50 ms, 250 ms, 500 ms and 1000 ms to narrow down the practical vitrification time limits. Consistent with our previous results difference density for the glucose molecule could be observed in the XI active site consistently across all time-points. This emphasizes that fast delay times are accessible to the *spitrobot* and that biologically relevant time-scales in the millisecond time-domain can be addressed via cryo-trapping crystallography.

**Supplementary Table 3:** Data collection and refinement statistics for CTX-M-14^E166A^ crystal datasets.

| CTX-M-14 |  |  |  |
| --- | --- | --- | --- |
| PDB-ID: | 8B2W | 8B2V | 8B2O |
| Delay time | 0.5 s | 1 s | 5 s |
| **Data collection** |  |  |  |
| Space group | P 31 2 1 | P 31 2 1 | P 31 2 1 |
| Cell dimensions (a, b, c (Å)) | 41.81 41.81 232.86 | 41.88 41.88 232.90 | 41.81 41.81 232.96 |
| $\alpha, \beta, \gamma$ (°) | 90.00 90.00 120.00 | 90.00 90.00 120.00 | 90.00 90.00 120.00 |
| Resolution (Å) |  | 77.64-1.73 | 77.65 - 1.86 |
| R merge | 0.22 (1.368) | 0.24 (1.193) | 0.27 (1.69) |
| I/$\sigma$(I) | 9.5 (2.0) | 9.9 (2.3) | 8.8 (2.0) |
| CC 1/2 | 99.9 (71.4) | 99.8 (72.4) | 99.8 (71.8) |
| Completeness (%) | 79.9 (32.2) | 80.2 (36.9) | 80.7 (37.2) |
| Redundancy | 10.1 (10.0) | 12.5 (10.0) | 15.3 (15.4) |
| Number of observations | 136189 (6755) | 181752 (7263) | 200593 (10125) |
| Number of unique | 13533 (677) | 14507 (726) | 13101(657) |
| **Refinement** |  |  |  |
| Resolution (Å) | 36.21 - 1.784 (1.848 - 1.784) | 38.82 - 1.73 (1.79 - 1.73) | 38.83 - 1.89 (1.96 - 1.89) |
| Total reflections | 27020 | 28951 | 26159 |
| Unique reflections | 13531 | 14503 | 13095 |
| R work / R free | 0.2328 / 0.2855 | 0.2009 / 0.2466 | 0.2142 / 0.2535 |
| **No. atoms** |  |  |  |
| Protein | 1967 | 2004 | 1962 |
| Ligand | 24 | 24 | 24 |
| Water | 171 | 188 | 146 |
| **B factors** |  |  |  |
| *Protein* | 21.79 | 22.1 | 27.39 |
| *Ligand* | 25.82 | 30.42 | 27.87 |
| *Water* | 25.61 | 30.15 | 34.94 |
| **R.m.s. deviations** |  |  |  |
| *Bond lengths (Å)* | 0.015 | 0.003 | 0.002 |
| *Bond angles (°)* | 1.76 | 0.67 | 0.56 |
| **Ramachandran statistics** |  |  |  |
| *Favored regions (%)* | 97.29 | 98.45 | 98.06 |
| *Allowed regions (%)* | 2.33 | 1.16 | 1.55 |
| *Outliers (%)* | 0.39 | 0.39 | 0.39 |

**Supplementary Table 4:** Data collection and refinement statistics for XI single crystal datasets.

|  | XI |  |  |  |
| --- | --- | --- | --- | --- |
| PDB-ID: | 8AWS | 8AWU | 8AWV | 8AWX |
| Delay time | 0.05 s | 0.25 s | 0.5 s | 1 s |
| **Data collection** |  |  |  |  |
| Space group | I 2 2 2 | I 2 2 2 | I 2 2 2 | I 2 2 2 |
| Cell dimensions (a, b, c (Å)) |  |  |  |  |
| $\alpha, \beta, \gamma$ (°) | 90.00 90.00<br>90.00 | 90.00 90.00<br>90.00 | 90.00 90.00<br>90.00 | 90.00 90.00<br>90.00 |
| Resolution (Å) | 68.99 - 2.263<br>(2.344 - 2.263) | 67.77 - 1.52<br>(1.574 - 1.52) | 51.19 - 1.58<br>(1.637 - 1.58) | 51.29 - 1.961<br>(2.031 - 1.961) |
| R merge | 0.273(2.11) | 0.713(245.7) | 0.144(1.433) | 0.126(1.039) |
| I/$\sigma$(I) | 8.6(1.4) | 10.5(1.32) | 10.6(1.6) | 11.5(2.1) |
| CC 1/2 | 99.3(55.9) | 99.0(30.9) | 99.8(74.8) | 99.8(74.8) |
| Completeness (%) | 84.3(51.1) | 99.08 (92.27) | 53.34 (5.35) | 85.64 (83.04) |
| Redundancy | 12.2(11.8) | 10.1(7.04) | 8.67(9.91) | 7.64(7.68) |
| Number of observations | 214819 |  | 297842 | 224568 |
| Number of unique | 17665 |  | 34368 | 29404 |
| **Refinement** |  |  |  |  |
| Resolution (Å) | 2.26 | 1.52 | 1.58 | 1.96 |
| Total reflections | 27111 | 733356 | 64529 | 40982 |
| Unique reflections | 17488 | 72319 | 34366 | 29381 |
| R work / R free | 0.1621/0.2233 | .01728/0.1889 | 0.15740.1946 | 0.1482/0.1917 |
| **No. atoms** |  |  |  |  |
| Protein | 3065 | 3131 | 3104 | 3173 |
| Ligand | 14 | 14 | 14 | 14 |
| Water | 238 | 522 | 389 | 376 |
| **B factors** |  |  |  |  |
| Protein | 23.75 | 13.73 | 21.05 | 23.52 |
| Ligand | 26.29 | 23.93 | 32.72 | 33.68 |
| Water | 27.23 | 28.66 | 32.95 | 34.66 |
| **R.m.s. deviations** |  |  |  |  |
| Bond lengths (Å) | 0.01 | 0.006 | 0.009 | 0.009 |
| Bond angles (°) | 1.12 | 0.91 | 1 | 1.02 |
| **Ramachandran statistics** |  |  |  |  |
| Favored regions (%) | 96.61 | 96.61 | 97.14 | 96.61 |
| Allowed regions (%) | 3.12 | 3.12 | 2.34 | 3.12 |
| Outliers (%) | 0.26 | 0.26 | 0.52 | 0.26 |

**Supplementary Figure 7:**
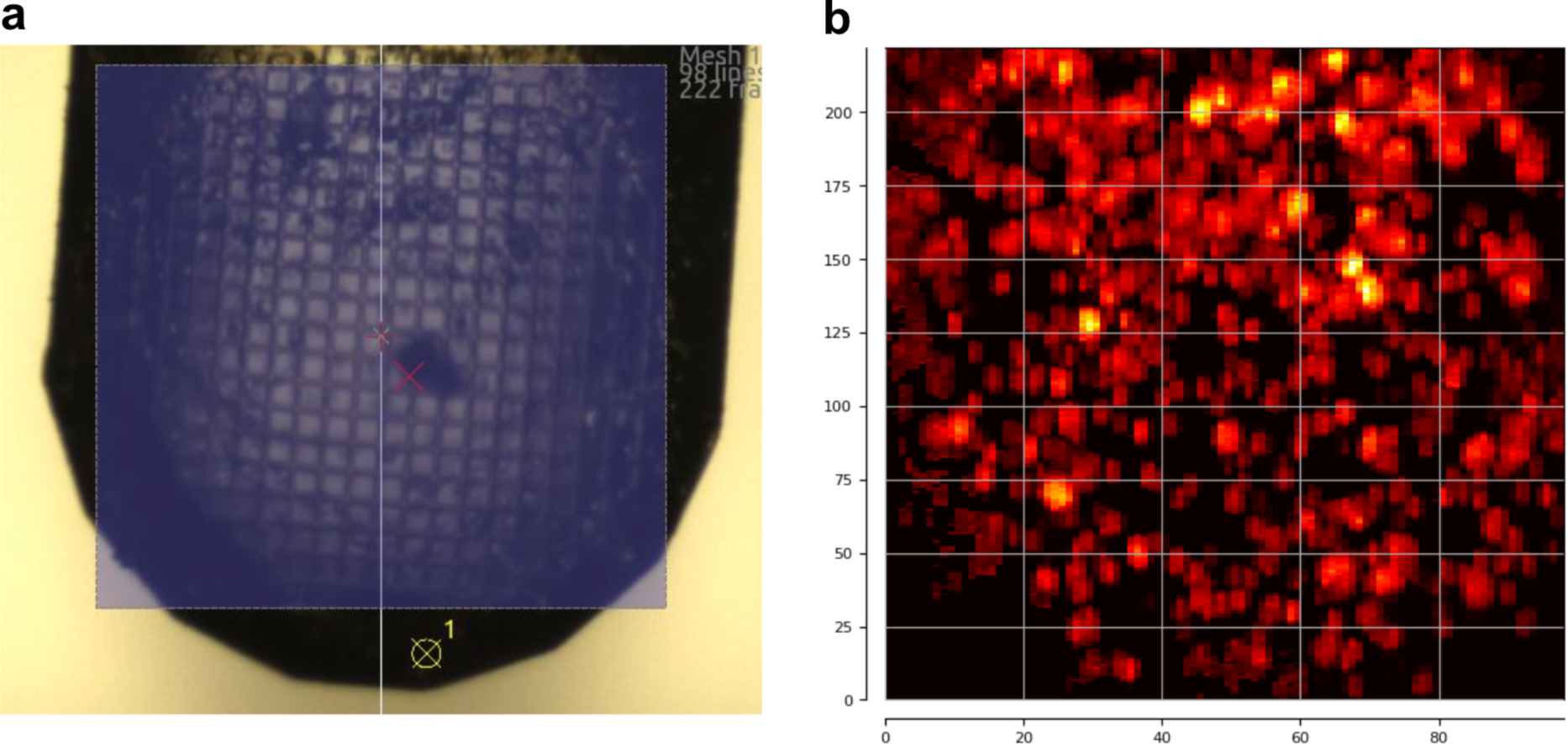
Mesh heat maps. The mesh approach can be utilized both in serial and single-crystal data collection. For cryo-SSX the heat map displays the diffraction quality of the crystals distributed on the mesh. For single-crystal data collection, the heat-maps help to identify the best diffracting crystals. a) an area on the micro-mesh is selected and raster scanned via a micro-focus X-ray beam. b) Diffraction quality results are displayed as a heat map on the same area, enabling convenient localization and selection of well-diffracting crystals. The heat map shows a typical distribution of micro-crystals on a micro-mesh.

### Cryo-trapping crystallography of tryptophan synthase reaction intermediates

Next, we aimed to demonstrate that the *spitrobot* can be used for time-resolved applications with macroscopic crystals. To this end we turned to tryptophan synthase (TS) as a pilot system. TS is a hetero-tetrameric pyridoxal 5’-phosphate (PLP) dependent bi-enzyme complex which generally assembles into a TrpA/TrpB_2_/TrpA. In this structural architecture, a ∼25 Å long allosteric communication tunnel connects each of the TrpA/TrpB heterodimers^14,41^. The accepted model for the TS turnover reaction is as following: TrpA reversibly converts indole-3-glycerol phosphate (IGP) into glyceraldehyde-3-phosphate (G3P) and indole. This is concomitant to a stabilization of the TrpA loop6. Simultaneously, in the active site of TrpB, serine reacts with PLP forming an external aldimine (Aex-Ser) intermediate. Tryptophan is finally generated by indole passing through the tunnel between TrpA^42^ and TrpB allosterically regulated by the communication domain (COMM) of the TrpB subunit (**Figure 3A**). Once indole reaches the TrpB active site it interacts with the Aex-Ser intermediate forming tryptophan as the end product, which is finally released from TS^41,43,44^.

To gain insight into reaction intermediates, TS macro-crystals were loaded onto standard 400/25 µm SPINE micromeshes (MiTeGen, USA) and mounted on the *spitrobot*. Turnover was initiated by LAMA-depositing reaction buffer containing indole, serine and glyceraldehyde-3-phosphate (G3P) as substrates onto the TS crystals. To this end 500 droplets of reaction buffer were deposited at a 6 kHz, repetition rate. The TS crystals were then vitrified in liquid nitrogen by plunging them directly into a puck at the specified delay time points (20 s, 25 s and 30 s).

At 20 and 30 seconds, in the active site of the TrpB subunit near full occupancy of the Aex-Ser intermediate are observed (**Figure 3**). In addition to the formation of this intermediate, the side chain movements of bLys87 and bGln114 are visible (**Supplementary Figure 8. a-d**).

These observations not only demonstrate that macroscopic crystals are amenable to the *spitrobot* but also that biochemical reaction intermediates can reliably be analyzed at delay times that would be inaccessible via manual cryo-trapping approaches.

**Supplementary Figure 8:**
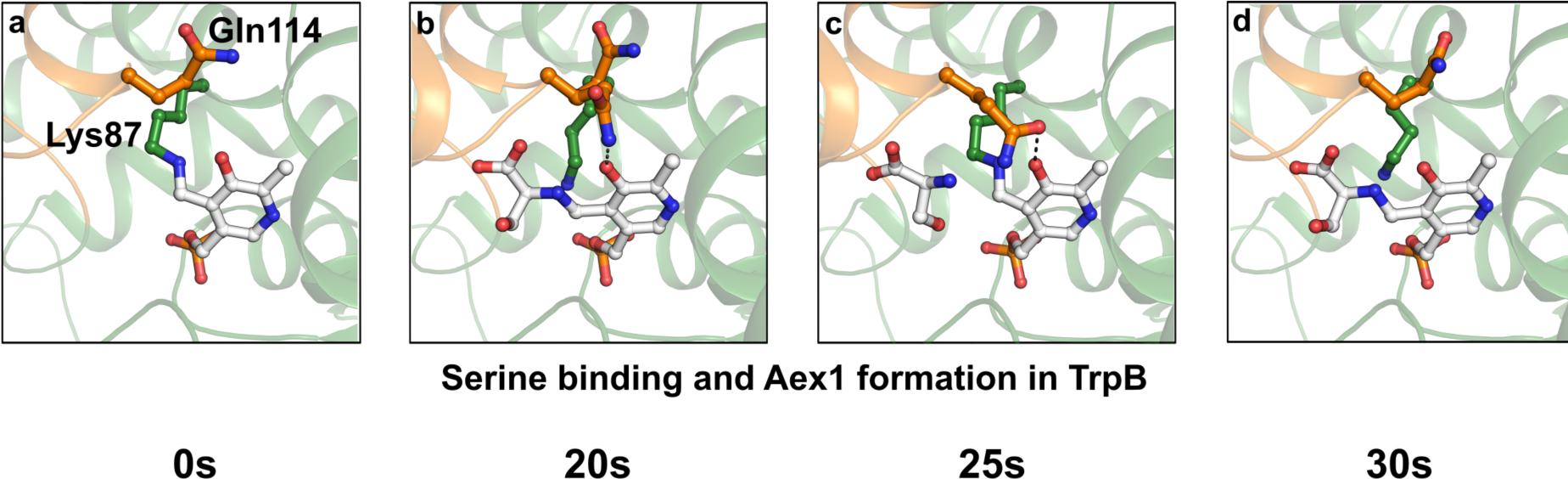
TS snapshots of the residues for serine binding and Aex1 formation in TrpB, after reaction initiation. **a-d)** Serine binding and Aex-Ser formation in TrpB Snapshots of residues involved in an external aldimine intermediate (Aex-Ser) formation in Trp B active site at 20 s and 30 s and serine binding state at 25 s after mixing. Active site residues of TrpB and substrates are represented as ball-and-stick.

**Supplementary Table 5:**
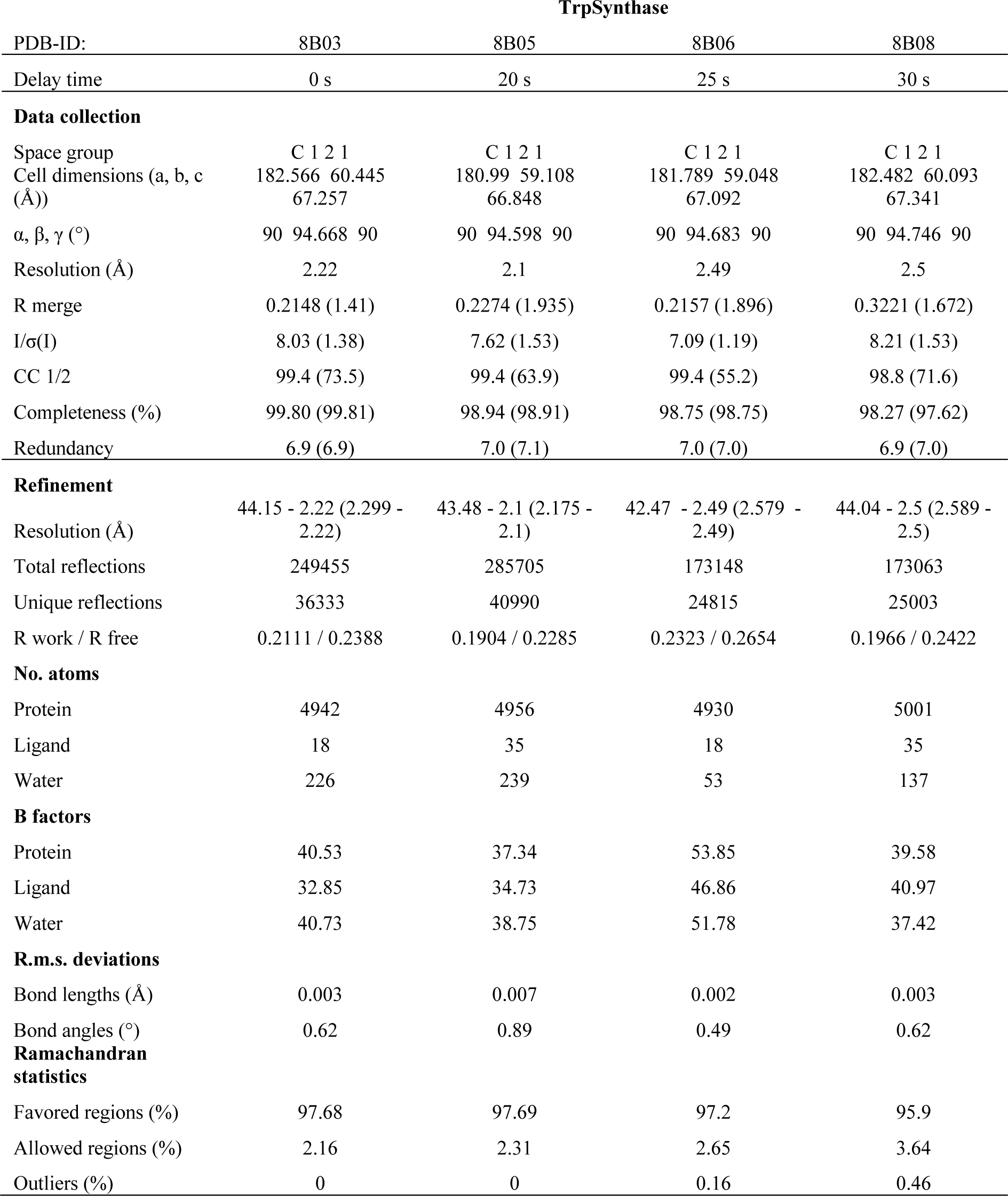
Data collection and refinement statistics for TS crystal datasets TrpSynthase.

